# Mass Cytometry Study on the Heterogeneity in Cellular Association and Cytotoxicity of Silver Nanoparticles in Human Immune Cells

**DOI:** 10.1101/617332

**Authors:** My Kieu Ha, Jang-Sik Choi, Zayakhuu Gerelkhuu, Sook Jin Kwon, Jaewoo Song, Yangsoon Lee, Yeoung-Eun Kim, Tae Hyun Yoon

## Abstract

There have been many reports about the adverse effects of nanoparticles (NPs) on the environment and human health. Conventional toxicity assessments of NPs frequently assume uniform distribution of monodisperse NPs in homogeneous cell populations, and provide information on the relationships between the administered dose of NPs and cellular responses averaged for a large number of cells. They may have limitations in describing the wide heterogeneity of cell-NP interactions, caused by cell-to-cell and NP-to-NP variances. To achieve more detailed insight into the heterogeneity of cell-NP interactions, it is essential to understand the cellular association and adverse effects of NPs at single-cell level. In this study, we applied mass cytometry to investigate the interactions between silver nanoparticles (AgNPs) and primary human immune cells. High dimensionality of mass cytometry allowed us to identify various immune cell types and observe the cellular association and toxicity of AgNPs in each population. Our findings showed that AgNPs had higher affinity with phagocytic cells like monocytes and dendritic cells and caused more severe toxic effects than with T cells, B cells and NK cells. Multi-element detection capability of mass cytometry also enabled us to simultaneously monitor cellular AgNP dose and intracellular signaling of individual cells, and subsequently investigate the dose-response relationships of each immune population at single-cell level, which are often hidden in conventional toxicity assays at bulk-cell level. Our study will assist future development of single-cell dose-response models for various NPs and will provide key information for the safe use of nanomaterials for biomedical applications.

## Introduction

As nanoparticles (NPs) have increasingly been employed in various consumer products and biomedical applications, their potential toxicity has become a public concern, since there have been many reports on the adverse effects of NPs on the environment and human health (Colvin 2003, Nel *et al.* 2006, Ray *et al.* 2009). It is important to examine the potential hazards of NPs on biological systems, especially on human immunity, which consists of various cell populations responsible for protecting our body from infections and malignancies. Conventional assays in nanotoxicity studies analyze the relationship between the administered dose of NPs and cellular responses, which are often averaged over a large number of cells. These assays at bulk-cell level may not able to describe cell-to-cell and NP-to-NP variances of cell-NP interactions under *in vivo* conditions (Singh *et al.* 2014). Knowledge of the biodistribution and toxicity of NPs in individual cells is essential for studying the dose-response relationship at a single-cell level and for gaining a more detailed understanding of cell-NP interactions.

To date, the biodistribution and toxicity of NPs in single cells have received noticeable attention but are mostly studied separately with only a few studies reporting on the single-cell dose-response relationship. On the one hand, a variety of quantitative and semiquantitative approaches have been employed to address the variability in the cellular uptake of NPs. Electron microscopy with a nanoscale resolution is suitable for examining the cellular localization of electron-dense NPs, but suffers from low-throughput and does not allow for the analysis of dynamic biological processes (Ivask, Mitchell, Malysheva, *et al.* 2017). Inductively coupled plasma mass spectrometry (ICP-MS) has been used to quantify the total metal concentration in a large number of cells after acid digestion, which is then averaged over the entire cell population to estimate the cellular NP content (Merrifield *et al.* 2018). This measurement can provide integrated information from thousands of cells, but will mask the stochastic diversity of individual cellular NP uptake (Meyer *et al.* 2018). New applications of ICP-MS based techniques have allowed for the single-cell quantification of cellular NP amounts. Laser ablation ICP-MS (LA-ICP-MS) enables the quantification of metal content in single cells with additional insights into subcellular localization. However, this image-based method suffers from low throughput and relatively low sensitivity (Yang *et al.* 2017). Single-particle ICP-MS utilizes a time-resolved mode to enable direct quantification of the concentration, size distribution, and agglomeration state of NPs (Yang *et al.* 2017). However, since it is not optimized for bioanalysis, it does not distinguish between cell-associated particles and unbound ones, so it should be coupled with other techniques with imaging capability such as LA-ICP-MS to determine the quantity and distribution of cellular NPs.

On the other hand, several techniques have also been applied to examine the *in vitro* toxicity of NPs, but few can be used to monitor both the cellular association and toxic effects of NPs simultaneously. Fluorescence-based microscopy is one of the few methods that can perform both tasks simultaneously, but it has low throughput and limited spatial resolution (200 – 500 nm) (Vanhecke *et al.* 2014). Moreover, due to the cross-talk and autofluorescence issues of the fluorophores, it fails to estimate the number of cell-associated NPs quantitatively (Orecchioni *et al.* 2017). Flow cytometry (FCM) is another technique that can simultaneously monitor cellular NP uptake and cellular responses by measuring the light-scattering and fluorescence-labeling intensities, respectively (Park *et al.* 2017, Ha *et al.* 2018). However, for those cells that have wide heterogeneity (e.g., immune cells), FCM is inadequate in providing simultaneous analyses of cell type, activation status, and elevation/depletion of secreted molecules due to its limited number of fluorescence detection channels. Recently, a combination of time-of-flight ICP-MS and FCM was developed and named as mass cytometry (Bandura *et al.* 2009, Bendall *et al.* 2011). It enables the detection of various metal-tagged cellular markers based on the mass-to-charge ratio of the elemental isotopes, with minimal overlap and cellular background interference. Current mass cytometry instruments allow up to 50 metal isotope labels (with atomic weights ranging from 75 to 209) to be detected simultaneously at single-cell resolution (Yang *et al.* 2017). This technique is suitable for a large panel design to analyze the cellular association and toxic effects of NPs in heterogeneous immune cells (Guo *et al.* 2017, Ivask, Mitchell, Hope, *et al.* 2017, Orecchioni *et al.* 2017).

Among the variety of currently available NPs, silver NPs (AgNPs) have been widely used for therapeutic interventions and medical diagnosis as drug carriers, nanoprobes, bio-imaging and labeling agents (Wei *et al.* 2015). At the same time, extensive studies have investigated the involvement of AgNPs in adverse biological effects, such as reactive oxygen species (ROS) generation, mitochondrial injury, DNA damage cell-cycle arrest and apoptosis induction (AshaRani *et al.* 2009, Ghosh *et al.* 2012, Martínez-Gutierrez *et al.* 2012, Ivask *et al.* 2014). AgNPs can enter the systemic circulation and cause cardiovascular toxicity due to their access to the heart, bloodstream and blood vessels (Takenaka *et al.* 2001, Nemmar *et al.* 2002). They have been found to exert toxicity on peripheral blood mononuclear cells (PBMCs), induce aggregation in platelets and promote procoagulant activity in red blood cells (Greulich *et al.* 2011, Jun *et al.* 2011, Bian *et al.* 2019).

In this study, we have adapted mass cytometry to investigate the cellular association and toxicity mechanisms of AgNPs in immune cells. Since the cellular association and cytotoxicity of AgNPs may be strongly influenced by the heterogeneous nature of PBMCs as well as the diverse physicochemical properties and colloidal behaviors of NPs, we have studied the impact of various cell types of PBMCs on the cellular association of AgNP, which led to even greater variances in signaling activities and apoptosis levels of individual cells with different phenotypes. Our findings will be useful in developing a single-cell dose-response model for future nanotoxicity studies based on the mass cytometry technique and will also provide information for the appropriate utilization of nanomaterials, especially AgNPs, in commercial and biomedical applications.

## Materials and methods

### Silver nanoparticles (AgNPs)

The AgNPs used in this study were BioPure Silver Nanospheres (nanoComposix, USA) with a polyvinylpyrrolidone coating and nominal core diameters of 10 and 20 nm (denoted ^PVP^Ag^10^ and ^PVP^Ag^20^, respectively).

### Isolation of peripheral blood mononuclear cells (PBMCs) from human whole blood

Whole blood was drawn from healthy donors into EDTA-treated tubes (BD Vacutainer®, USA), and PBMCs were isolated from whole blood via density gradient centrifugation using Ficoll-Paque PLUS (GE Healthcare Bio-Sciences, Sweden). Briefly, blood was diluted 1:1 with Dulbecco’s phosphate-buffered saline (DPBS, Welgene, Korea), and the diluted blood was overlaid on the Ficoll reagent in centrifuge tubes. These tubes were centrifuged at 400 × *g*, at room temperature for 40 minutes on a centrifuge with a swing-bucket rotor (model 1248R, Labogene, Korea). The mononuclear layer was then collected and transferred to a new tube, washed in DPBS, and pelleted by centrifugation at 200 × *g*, at room temperature for 10 minutes. The resulting supernatant was discarded, and the cells were resuspended in RPMI-1640 medium (Lonza™ BioWhittaker™, USA) supplemented with 10% fetal bovine serum (Gibco, USA) and 1% penicillin/streptomycin (Gibco, USA). Cell suspensions were transferred to 35 × 10 mm petri dishes (SPL Life Sciences, Korea) for subsequent treatment with AgNPs.

### Sample preparation for mass cytometry measurements

Cells were incubated with 2 or 5 μg/mL of ^PVP^Ag^10^ or ^PVP^Ag^20^ NPs in RPMI complete media for 3 h at 37°C and 5% CO_2_. Then, cells were labelled with surface and intracellular markers according to the Maxpar Phosphoprotein Staining with Fresh Fix Protocol (Fluidigm Corp., USA). Briefly, cells were washed with DPBS to remove excess AgNPs and then stained with Cisplatin for viability. After that, cells were stained with surface antibodies (Table 1). After surface staining, cells were fixed in 1.6% formaldehyde and permeabilized with ice-cold methanol. Then, cells were stained with phosphoprotein antibodies (Table 1). After intracellular staining, cells were incubated with Cell-ID Intercalator-Ir solution overnight. Prior to data acquisition, cells were washed and suspended at 1 × 10^6^ cells/mL in ultrapure water. Calibration beads were added 1:10 v/v for later normalization. Cells were then filtered into strainer cap tubes and ready for acquisition.

**Table 1.** List of antibodies in the labeling panel.

| Surface antibodies |  |  |  |  |  |
| --- | --- | --- | --- | --- | --- |
| Target | Clone | Metal | Target | Clone | Metal |
| CD11a | HI111 | <sup>142</sup>Nd | CD197 (CCR7) | G043H7 | <sup>159</sup>Tb |
| CD4 | RPA-T4 | <sup>145</sup>Nd | CD45RO | UCHL1 | <sup>165</sup>Ho |
| CD8a | RPA-T8 | <sup>146</sup>Nd | CD44 | BJ18 | <sup>166</sup>Er |
| CD16 | 3G8 | <sup>148</sup>Nd | CD45RA | HI100 | <sup>169</sup>Tm |
| CD7 | CD7-6B7 | <sup>147</sup>Sm | CD3 | UCHT1 | <sup>170</sup>Er |
| CD45 | HI30 | <sup>154</sup>Sm | HLA-DR | L243 | <sup>174</sup>Yb |
| Intracellular protein antibodies |  |  |  |  |  |
| Target | Clone | Metal | Target | Clone | Metal |
| pSTAT5 | 47 | <sup>150</sup>Nd | pBAD | 40A9 | <sup>161</sup>Dy |
| Caspase-7 | D6H1 | <sup>152</sup>Sm | IκBα | L35A5 | <sup>164</sup>Dy |
| pSTAT1 | 58D6 | <sup>153</sup>Eu | pERK1/2 | D13.14.4E | <sup>171</sup>Yb |
| p38 | D3F6 | <sup>156</sup>Gd | pS6 | N7-548 | <sup>175</sup>Lu |
| pSTAT3 | 4/P-Stat3 | <sup>158</sup>Gd |  |  |  |

### Data acquisition and analysis

The samples were analyzed on the Helios mass cytometry platform (Fluidigm Corp., USA) at a flow rate of 45 μL/min, in dual instrument mode with noise reduction turned on, and with ‘on the fly’ processing. Measurements were performed in triplicate. The initial raw data were processed into standard FCS format and normalized by the CyTOF software version 6.5.358 (Fluidigm Corp., USA).

FlowJo (FlowJo, LLC, USA) and Cytobank (Cytobank, Inc., USA) were used for data gating and visualization. Inverse hyperbolic sine (arcsinh) transformation was applied to the raw data. The major immune cell subsets were then gated using the 12 surface markers (Table 1). The gating pipeline is presented in Figure S1. The SPADE (spanning tree progression analysis of density-normalized events) clustering algorithm was used to reduce the dimensionality of the data. From the SPADE plots, we identified the cell subsets that had high cellular AgNP uptake and cell death levels based on the intensities of ^107^Ag and ^195^Pt. Next, the median expression levels of 9 phosphoprotein markers (Table 1) were plotted on heatmaps in order to understand the signaling pathways of AgNP-induced toxicity. Then, the t-distributed stochastic neighbor embedding (t-SNE) dimension-reduction method was used to visualize single-cell resolution of the data. The bivariate plots between cellular responses and AgNP uptake were used to analyze the dose-response relationship at a single-cell level.

Quantification of cellular AgNP uptake was done using Ag^+^ ion calibration, in which a standard Ag^+^ ion solution diluted in 5% HNO_3_ was analyzed in “solution mode”. The transmission efficiency (TE) of a given number of ^107^Ag atoms was determined as in equations 1 and 2 (Tricot *et al.* 2015).

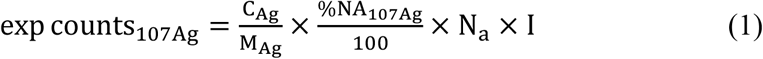

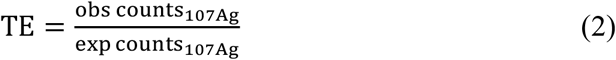

where C_Ag_ and obs counts_107Ag_ are the concentration (μg/mL) and observed ^107^Ag counts from the standard Ag^+^ ion solution, M_Ag_ is the atomic mass of Ag, %NA_107Ag_ is the natural abundance percentage of ^107^Ag (51.839%), N_a_ is Avogadro’s number (6.02 × 10^23^), and I is the injection speed (or flow rate).

Cell samples were analyzed in “event mode” and their expected cell-associated ^107^Ag signal (exp cell_107Ag_) was calculated from the observed ^107^Ag signal (obs cell_107Ag_) and TE as shown in equation 3. Then, the particle mass and number of cell-associated Ag could be calculated using equation 4 (Ivask, Mitchell, Hope, *et al.* 2017).

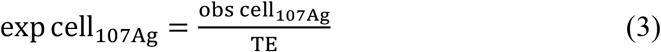

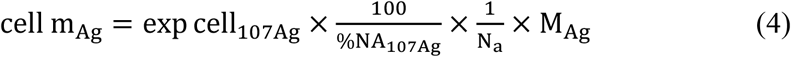

## Results

### Cellular uptake of AgNPs

To identify the major immune cell subsets, we applied the SPADE clustering algorithm (Bendall *et al.* 2011) to the 12 surface markers (Table 1). This approach uses a minimum-spanning tree algorithm, in which each tree node encompasses a cluster of cells that are phenotypically similar in the 12-dimensional space defined by the surface markers, and the connections between nodes display the relationships between the cell clusters. The SPADE trees provide a convenient approach to map complex, high-dimensional data into a representative two-dimensional structure. The intensity of ^107^Ag represents the cellular uptake of AgNPs. The SPADE trees show that cellular uptake of AgNPs was not evenly distributed in all populations (Figure 1). The uptake amounts of ^PVP^Ag^10^ and ^PVP^Ag^20^ NPs were relatively similar, with that of ^PVP^Ag^20^ NPs (Figure 1B,C,D) being slightly higher than that of ^PVP^Ag^10^ NPs (Figure 1E,F,G). The 2 types of AgNPs only had subtle difference in cellular uptake amounts because the difference in their sizes was relatively small (only 2 fold). Among the identified populations, monocytes and dendritic cells generally had higher AgNP uptake amounts than other cell types. Because they are phagocytic cells, monocytes and dendritic cells have a higher tendency to ingest foreign particles than other non-phagocytic cells (Zolnik *et al.* 2010, Luo *et al.* 2015).

**Figure 1.**
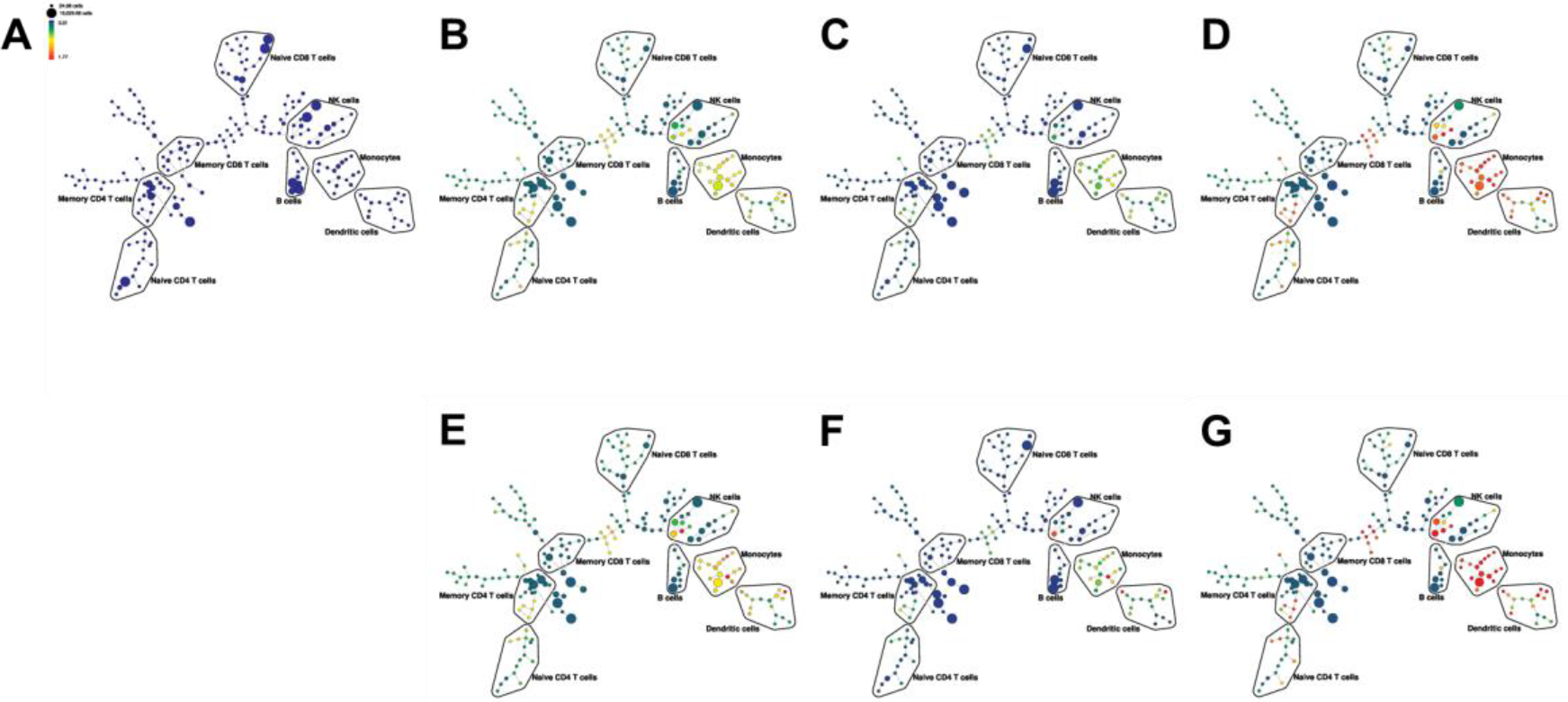
SPADE plots of cellular AgNP uptake in (A) control and (B-G) AgNP-treated samples. Exposure conditions were (B) ^PVP^Ag^10^ NPs at 2 μg/mL for 3 h, (C) ^PVP^Ag^10^ NPs at 2 μg/mL for 6 h, (D) ^PVP^Ag^10^ NPs at 5 μg/mL for 3 h, (E) ^PVP^Ag^20^ NPs at 2 μg/mL for 3 h, (F) ^PVP^Ag^20^ NPs at 2 μg/mL for 6 h, and (G) ^PVP^Ag^20^ NPs at 5 μg/mL for 3 h. Node color is scaled to the median mass of cell-associated Ag (in fg), with the lighter color (e.g. red) representing high intensity and the darker color (e.g. blue) representing low intensity.

In the samples treated for the same exposure time (3 h) but with different doses of AgNP (2 vs. 5 μg/mL) (Figure 1B and 1D, Figure 1E and 1G), the cellular AgNP content increased as the administered dose increased. This agrees with our previous study regarding the dose-dependent cellular uptake of AgNPs (Ha *et al.* 2018).

However, when comparing samples treated with the same AgNP dose (2 μg/mL) but for different exposure time (3 vs. 6 h) (Figure 1B and 1C, Figure 1E and 1F), cellular AgNP content decreased as the exposure time increased, most obviously in monocytes and dendritic cells. This observation suggests that after 6 h, the phagocytosed AgNPs were either broken down or excreted, making their cellular dose decrease.

### Cellular response to AgNP exposure

Figure 2 displays the cell death levels of identified populations after AgNP treatment. The cell death level is represented by the intensity of cisplatin, which is a molecule that can penetrate late apoptotic and necrotic cells that have lost their membrane integrity. As shown in Figure 2, the cell death levels were different between populations. Monocytes and dendritic cells had the highest cell death levels among all identified populations, which corresponded to their high AgNP uptake. Their cell death levels also decreased when the exposure time increased from 3 h to 6 h but increased when the administered dose increased from 2 μg/mL to 5 μg/mL. The corresponding patterns in cellular AgNP uptake and cell death levels confirmed that the cell death in the identified immune populations was caused by the cellular association and toxic effects of AgNPs.

**Figure 2.**
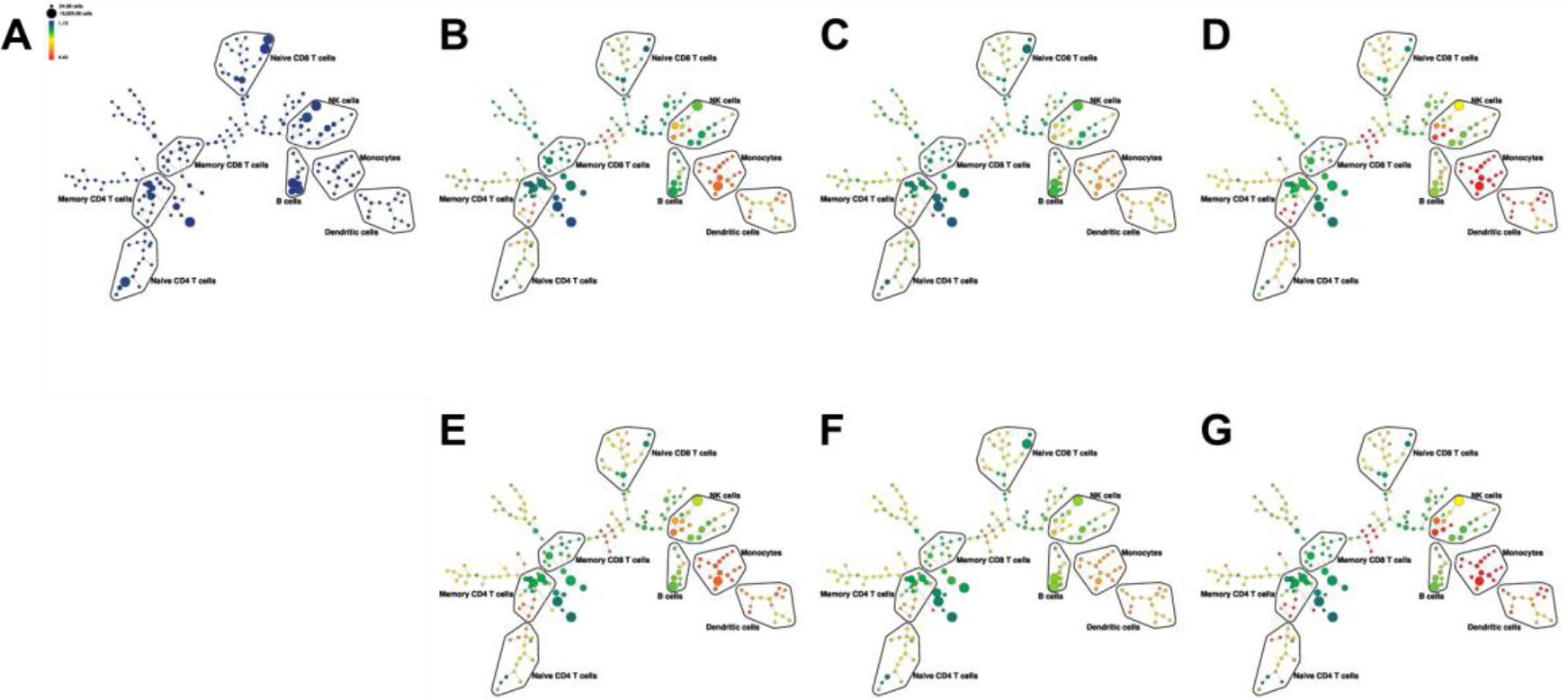
SPADE plots of cell death in (A) control and (B–G) AgNP-treated samples. Exposure conditions were (B) ^PVP^Ag^10^ NPs at 2 μg/mL for 3 h, (C) ^PVP^Ag^10^ NPs at 2 μg/mL for 6 h, (D) ^PVP^Ag^10^ NPs at 5 μg/mL for 3 h, (E) ^PVP^Ag^20^ NPs at 2 μg/mL for 3 h, (F) ^PVP^Ag^20^ NPs at 2 μg/mL for 6 h, and (G) ^PVP^Ag^20^ NPs at 5 μg/mL for 3 h. Node color is scaled to the median intensity of cisplatin expression, with the lighter color (e.g. red) representing high intensity and the darker color (e.g. blue) representing low intensity.

These observations suggest that a short-exposure at a high dose of AgNPs will result in higher cellular association and eventually higher toxicity compared to long-exposure at a low dose, especially in monocytes and dendritic cells. This means that in phagocytic cells like monocytes and dendritic cells, AgNPs can be ingested and cause toxic effects more quickly and efficiently than in other non-phagocytic cells. Therefore, it is important to understand by which signaling pathways the AgNPs exert their toxic effects and induce cell death.

In order to understand the signaling pathways behind AgNP-induced toxicity, we used heatmaps to visualize the median intensity of 9 intracellular protein markers in the identified immune populations. Figure 3 shows that the toxic effects of AgNPs did not alter the expression levels of all 9 signaling proteins, but only those of IκBα, STAT1, and caspase 7. However, at such low doses of AgNPs (i.e. 2 and 5 μg/mL), the expression levels of IκBα, STAT1, and caspase 7 in T cells, B cells, and NK cells in AgNP-treated samples were not significantly different from those of control. Only in monocytes and dendritic cells, the differential expressions of these signaling proteins were found especially significant. Compared to the control, in AgNP-treated samples, IκBα was downregulated while STAT1 and cleaved caspase 7 were upregulated in monocytes and dendritic cells. This finding suggests that exposure to AgNPs led to the activation of contrasting and complex cellular signaling pathways: STAT1 phosphorylation to promote inflammation, NF-κB signaling (via degradation of IκBα) to prevent apoptosis, and activation of caspase 7 to cleave downstream substrates that can eventually lead to apoptosis. It also indicates that low doses of NPs (in this case, 5 μg/mL or less) may cause weak and chronic adverse effects in the signaling functionality of T cells, B cells, and NK cells, but strong and acute toxicity in monocytes and dendritic cells.

**Figure 3.**
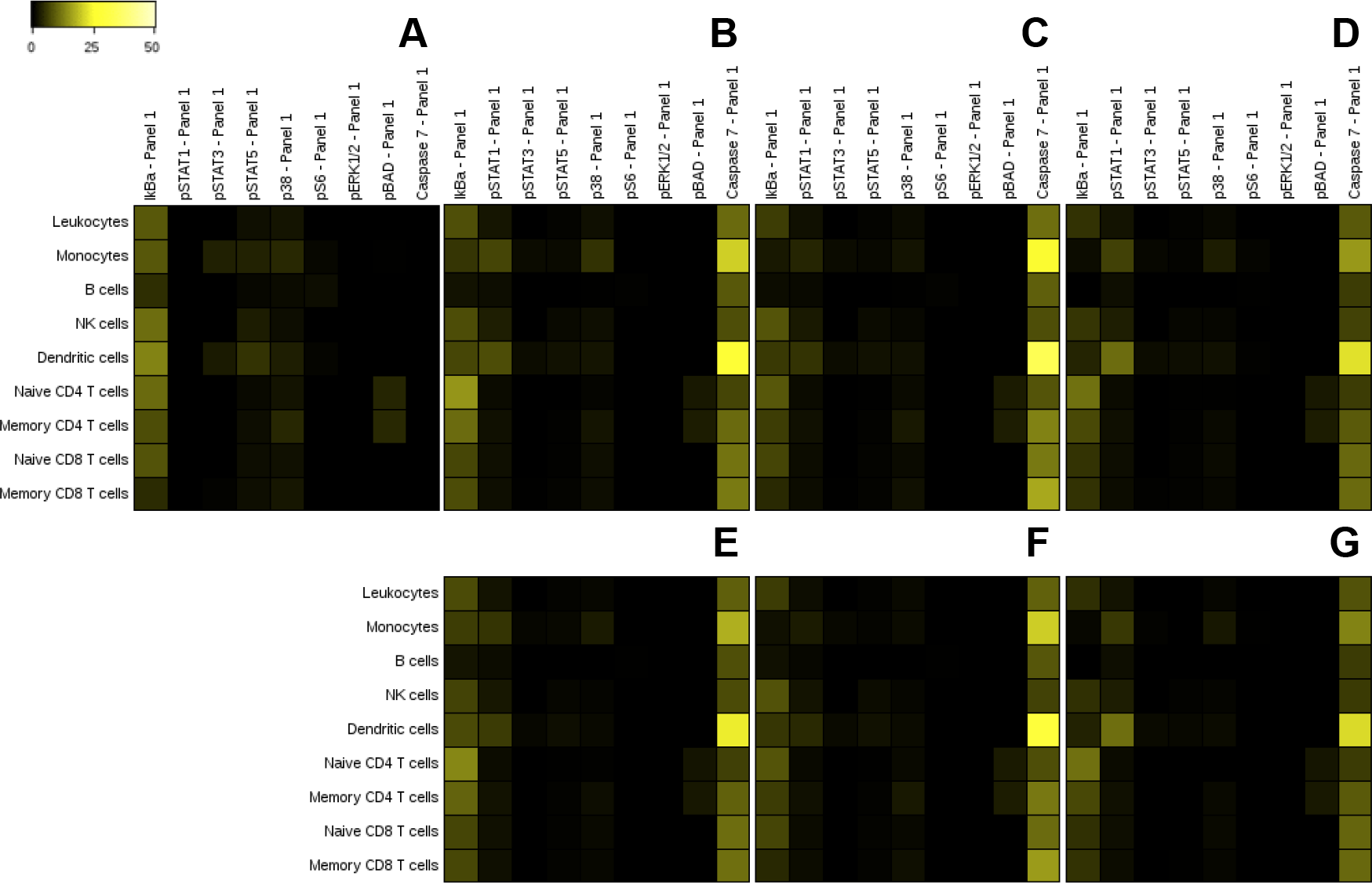
Heatmaps of phosphoprotein markers’ expression levels in (A) control and (B–G) AgNP-treated samples. Exposure conditions were (B) ^PVP^Ag^10^ NPs at 2 μg/mL for 3 h, (C) ^PVP^Ag^10^ NPs at 2 μg/mL for 6 h, (D) ^PVP^Ag^10^ NPs at 5 μg/mL for 3 h, (E) ^PVP^Ag^20^ NPs at 2 μg/mL for 3 h, (F) ^PVP^Ag^20^ NPs at 2 μg/mL for 6 h, and (G) ^PVP^Ag^20^ NPs at 5 μg/mL for 3 h. The color scale indicates the median intensity of each marker.

Despite providing simple and easy-to-interpret visualization, SPADE plots and heatmaps do not preserve the single-cell resolution of mass cytometry data. To make a representative visualization of cellular AgNP uptake and cell death as well as the induction of the differentially-expressed signaling proteins (i.e. IκBα, STAT1, and caspase 7) with single-cell resolution, we applied the t-SNE dimension reduction method, which is a computational approach for visualization of high-dimensional data with single-cell resolution (Van Der Maaten and Hinton 2008). t-SNE clustered the single cells into populations according to similarities in the multi-dimensional phenotypic expression of 12 surface proteins (Table 1) and projected the clusters onto bi-axial plots, as in Figure 4 (Characterization of each cell type in each sample is presented in Figure S3-S10).

**Figure 4.**
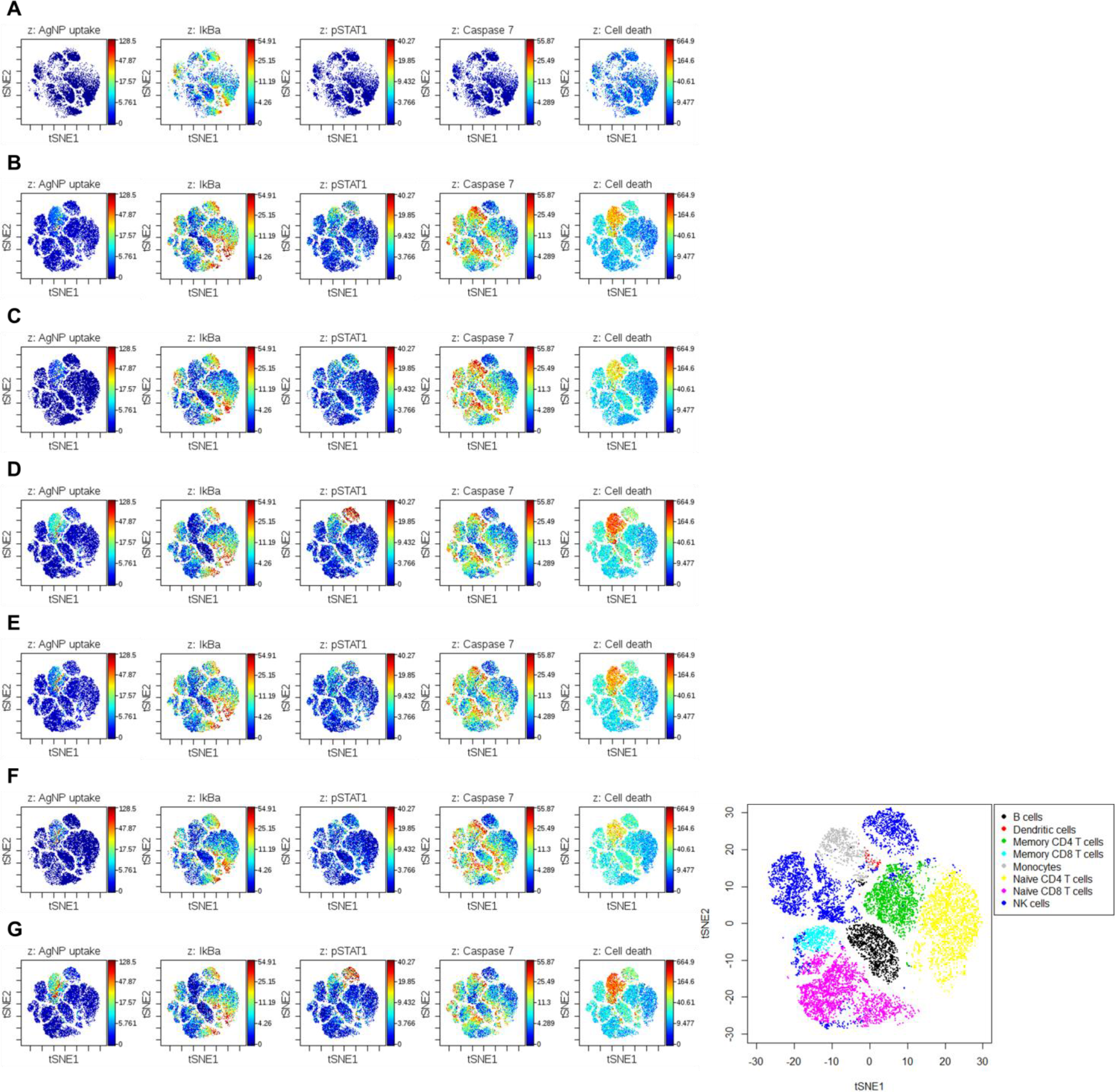
t-SNE plots of cellular AgNP uptake (in fg) and IκBα, STAT1, caspase 7, and cell death levels in (A) control and (B–G) AgNP-treated samples. Exposure conditions were (B) ^PVP^Ag^10^ NPs at 2 μg/mL for 3 h, (C) ^PVP^Ag^10^ NPs at 2 μg/mL for 6 h, (D) ^PVP^Ag^10^ NPs at 5 μg/mL for 3 h, (E) ^PVP^Ag^20^ NPs at 2 μg/mL for 3 h, (F) ^PVP^Ag^20^ NPs at 2 μg/mL for 6 h, and (G) ^PVP^Ag^20^ NPs at 5 μg/mL for 3 h. The color gradient represents the intensity of marker expression, in which the lighter color (e.g. red) shows high intensity and the darker color (e.g. blue) shows low intensity. Plot scale and cell type labeling are displayed at the bottom right corner.

The single-cell resolution obtained with the t-SNE analysis demonstrated heterogeneity in cellular AgNP uptake and the signaling expression profiles of PBMCs after AgNP-treatment. t-SNE results supported the cellular AgNP uptake and toxicity data observed in the SPADE plots. These results show that cellular uptake of ^PVP^Ag^10^ and ^PVP^Ag^20^ NPs occurred quickly (within 3 h) in monocytes and dendritic cells, exerting remarkable toxicity; conversely, uptake was slower in T cells, B cells, and NK cells, and did not cause severe cell death. They also show that AgNP uptake and cell death levels slightly decreased when the exposure duration was extended from 3 h to 6 h but increased when the exposure dose increased from 2 μg/mL to 5 μg/mL. The expression levels of IκBα, STAT1, and caspase 7 signaling markers in t-SNE plots also supported the heatmap findings: compared to control cells, IκBα was downregulated while STAT1 and caspase 7 were upregulated in samples exposed to AgNPs. However, the differential expression levels of IκBα, STAT1, and caspase 7, as well as AgNP uptake and cell death levels did not vary much between samples treated with ^PVP^Ag^10^ and ^PVP^Ag^20^ NPs. This indicates that the 2-fold size difference did not cause any notable changes in the toxic effects of these two NPs.

### Dose-response relationship at a single-cell level

Because monocytes and dendritic cells were two populations that showed more significant AgNP uptake and cellular responses than the other cell types, we wanted to investigate their dose-response relationships in more detail. The scatter plots of IκBα, STAT1, caspase 7, and cell death levels versus the AgNP uptake of monocytes and dendritic cells in Figures 5 and 6 confirm the previous observations about IκBα, STAT1, caspase 7, and cell death levels from Figures 1–4 in a single-cell dose-dependent manner.

**Figure 5.**
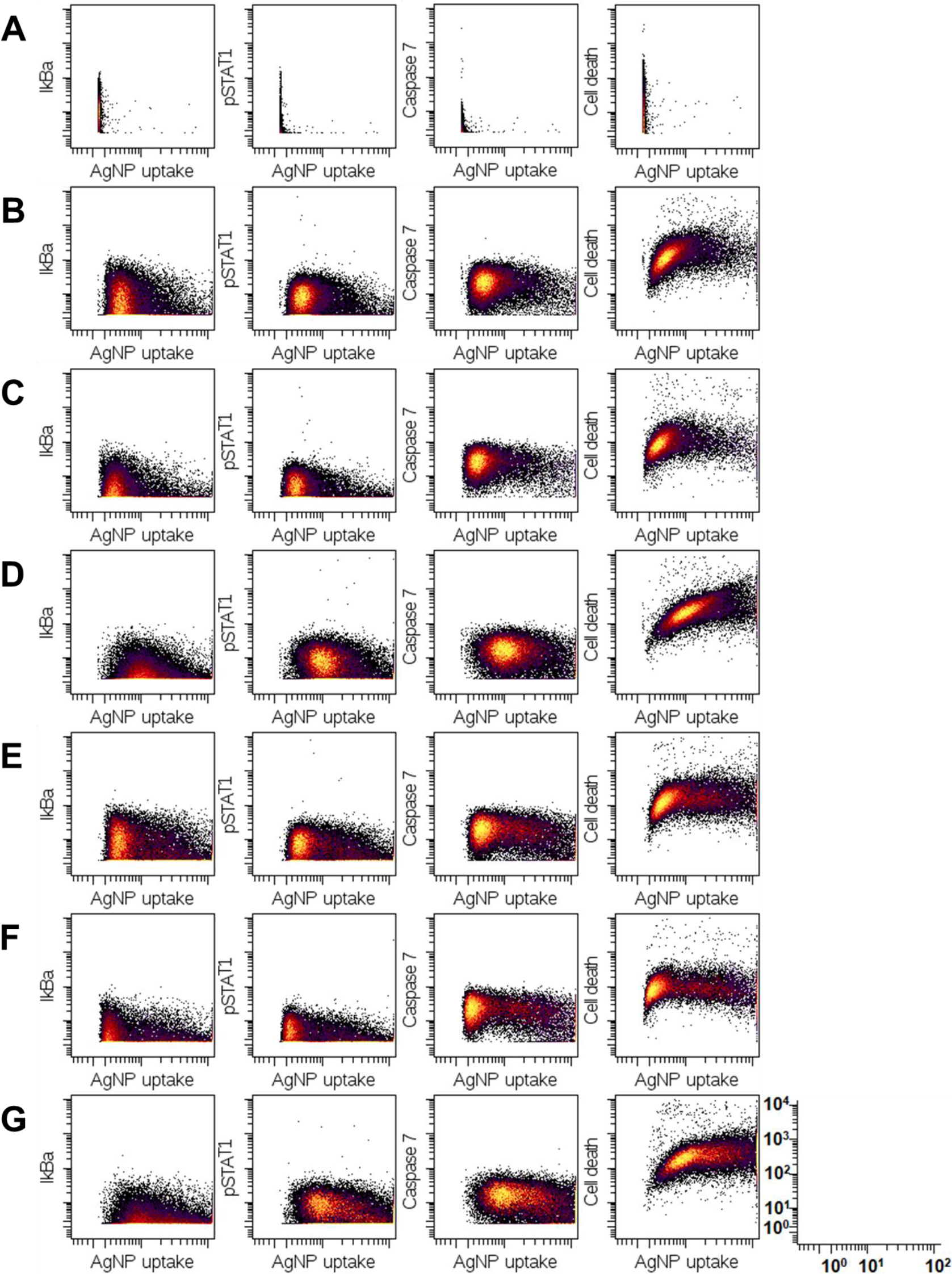
Scatter plots of IκBα, STAT1, caspase 7, and cell death levels versus AgNP uptake (in fg) of monocytes in (A) control and (B–G) AgNP-treated samples. Exposure conditions were (B) ^PVP^Ag^10^ NPs at 2 μg/mL for 3 h, (C) ^PVP^Ag^10^NPs at 2 μg/mL for 6 h, (D) ^PVP^Ag^10^ NPs at 5 μg/mL for 3 h, (E) ^PVP^Ag^20^ NPs at 2 μg/mL for 3 h, (F) ^PVP^Ag^20^ NPs at 2 μg/mL for 6 h, and (G) ^PVP^Ag^20^ NPs at 5 μg/mL for 3 h. The color gradient represents cell density, in which the lighter color (e.g. yellow) shows high density and the darker color (e.g. black) shows low density. The scale is displayed at the bottom right corner.

**Figure 6.**
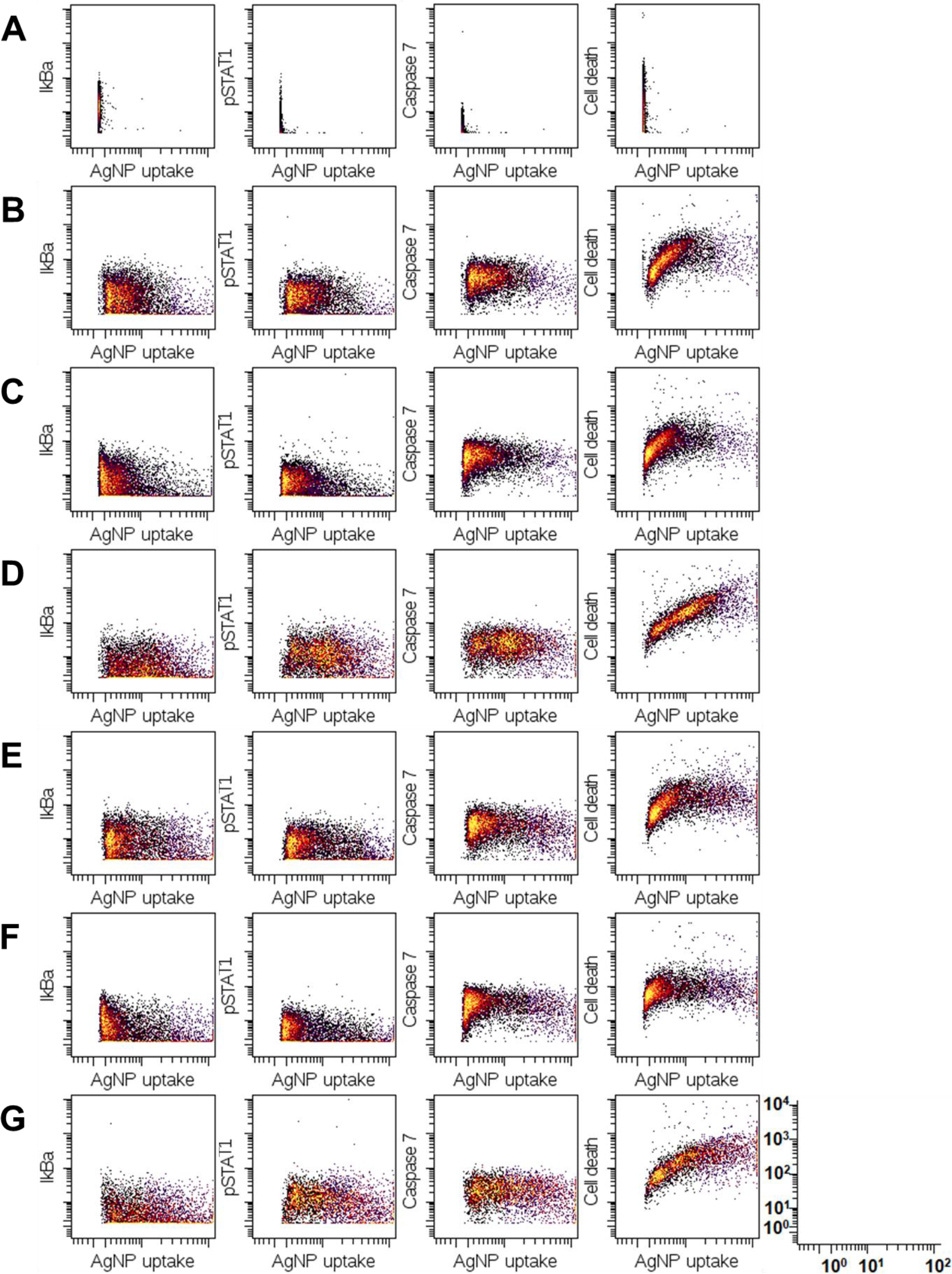
Scatter plots of IκBα, STAT1, caspase 7, and cell death levels versus AgNP uptake (in fg) of dendritic cells in (A) control and (B-G) AgNP-treated samples. Exposure conditions were (B) ^PVP^Ag^10^ NPs at 2 μg/mL for 3 h, (C) ^PVP^Ag^10^ NPs at 2 μg/mL for 6 h, (D) ^PVP^Ag^10^ NPs at 5 μg/mL for 3 h, (E) ^PVP^Ag^20^ NPs at 2 μg/mL for 3 h, (F) ^PVP^Ag^20^ NPs at 2 μg/mL for 6 h, and (G) ^PVP^Ag^20^ NPs at 5 μg/mL for 3 h. The color gradient represents cell density, in which the lighter color (e.g. yellow) shows high density and the darker color (e.g. black) shows low density. The scale is displayed at the bottom right corner.

The dose-response relationships in monocytes and dendritic cells were similar. Through observation at single-cell resolution, we can see that the cellular uptake of ^PVP^Ag^20^ NPs was in fact slightly higher than that of ^PVP^Ag^10^ NPs, as there were more cells with over 10 fg of AgNP uptake in the ^PVP^Ag^20^-treated sample than in the ^PVP^Ag^10^-treated sample. The degradation of IκBα and the phosphorylation of STAT1 were similarly dependent on cellular AgNP uptake, since IκBα and phospho-STAT1 levels were both inversely proportional to the AgNP uptake amounts. Cleaved caspase 7 levels in AgNP-treated samples was higher than that of the control but did not display clear dose-dependency. Unlike the other 3 markers mentioned above, cell death levels displayed a direct linear relationship with cellular AgNP uptake. This relationship maintained its linearity in the ^PVP^Ag^10^-treated samples (Figures 5,6B-D), as the cell death level increased accordingly with increases in cellular Ag dose, from mostly below 10 fg (Figures 5,6B and 5,6C) to over 10 fg, and even reaching 100 fg (Figures 5,6D). However, this linear relationship was not very consistent in the ^PVP^Ag^20^-treated samples (Figures 5,6E-G). In these samples, linearity was only maintained when the uptake amounts were below 10 fg. Above this threshold, the uptake amounts reached a plateau and did not increase any further.

## Discussion

Mass cytometry allowed us to simultaneously analyze 21 markers to differentiate distinct populations of PBMCs and to examine their differential signaling expressions with high sensitivity and throughput in order to understand the immune response to nanomaterial exposure.

As silver NPs (AgNPs) have been increasingly used in consumer products and biomedical applications due to their antimicrobial properties, they have also been reported to enter the systemic circulation and exert toxicity on PBMCs, induce aggregation in platelets, and promote procoagulant activity in red blood cells (Greulich *et al.* 2011, Jun *et al.* 2011, Bian *et al.* 2019). When NPs come into contact with immune cells, they are often first taken up by the phagocytic cells (e.g. monocytes, macrophages, and dendritic cells) (Zolnik *et al.* 2010). However, internalization of NPs can also occur in other immune cells by lysosomal or endosomal endocytosis. The compatibility of NPs with the immune system is largely determined by their physicochemical properties, such as size and surface chemistry (Dobrovolskaia and McNeil 2007). When NPs are taken up by immune cells, there are several possibilities for how they will interact with one another: the first scenario is immune-mediated destruction or rejection, which could initiate a defensive immune reaction resulting in elimination of the NPs; the second is immunostimulation, which promotes inflammatory or autoimmune disorders; and the third is immunosuppression, which increases the host’s susceptibility to infections (Luo *et al.* 2015). Oxidative stress and the release of metal ions have been reported as the two primary mechanisms that mediate the cytotoxicity of AgNPs (Völker *et al.* 2013). The acidic environment in lysosomes—where AgNPs are abundantly accumulated—stimulates the intracellular release of metal ions. These ions and the cell-associated AgNPs may escape into the cytosol and subsequently cause oxidative stress via a ROS-dependent pathway. On the other hand, in a ROS-independent manner, AgNPs and the released Ag^+^ ions may interact with the thiol groups of proteins, resulting in protein misfolding and eventually cell death (Riaz Ahmed *et al.* 2017). Metal ions have also been confirmed to mediate the toxicological pathways of metal-bearing NPs in a multiomics study that involved direct infusion mass spectrometry lipidomics, polar metabolomics, and RNA-sequencing transcriptomics (Dekkers *et al.* 2018). In this study, the majority of the molecular responses of human lung epithelial A549 cells to silver (Ag) and zinc oxide (ZnO) NPs—such as metallothionein induction, antioxidant depletion, repressed DNA repair, and apoptosis induction—are similar to their responses to Ag^+^ and Zn^2+^ ions, respectively, confirming that the modes of action of these NPs are largely mediated by the corresponding dissolved metal ions.

Low concentrations (2 and 5 μg/mL) of AgNPs were chosen for our experiments to avoid the saturation of silver intensity on the Helios platform and to maintain adequate cellular viability (Ivask *et al.* 2016), while still ensuring noticeable changes in cells’ viability and signaling functions. Our results demonstrate that monocytes and dendritic cells had more affinity for AgNPs and were thus more vulnerable to their toxicity. From the cellular Ag content shown in the SPADE plots (Figure 1) and t-SNE plots (Figure 3), we can see that within 3 to 6 h of exposure, the cellular AgNP content in these cells was higher than in other populations. This cell-type-specific AgNP uptake can be explained by the intrinsic properties and functions of the cells. Monocytes and dendritic cells are partly adherent cells, so they can interact with both suspended and sedimented AgNPs in culture media. Furthermore, these cells belong to the cytic cell group, so they naturally tend to phagocytose the administered AgNPs (Luo *et al.* 2015). T cells and B cells, on the other hand, are non-adherent cells, so they can only interact with suspended AgNPs. Moreover, one of the functions of these cells is to generate cellular responses to remove foreign pathogens and particles (Luo *et al.* 2015), so they tend to eliminate AgNPs rather than phagocytose them.

Exposure to AgNPs led to activation of contrasting and complex cellular signaling pathways: STAT1 phosphorylation, IκBα degradation, and caspase 7 activation. IκBα is a molecule that can inhibit the anti-apoptotic NF-κB transcription factor; therefore, degradation of IκBα by phosphorylation can induce NF-κB transcriptional activity and thus prevent apoptosis (Viatour *et al.* 2005). STAT1 is a transcription factor in the STAT protein family, which is a critical part of the JAK-STAT signaling pathway. Phosphorylation of STAT1 can prevent proliferation, promote inflammation, and induce apoptosis (Schindler *et al.* 2007). Caspase 7 belongs to the apoptosis executioner protein family, which is activated through proteolytic processing at conserved aspartic residues by upstream enzymes, including caspase 3, 6, 8, and 9, along with granzyme B. Upon activation, it will cleave downstream substrates and eventually lead to apoptosis (Lamkanfi and Kanneganti 2010). The contrasting expression patterns of the aforementioned pathways reflect the cells’ efforts to achieve a more balanced response to the toxic effects of AgNP exposure.

The differences in phosphoprotein signaling and viability between cells treated with 2 μg/mL and those treated with 5 μg/mL of AgNPs were caused by the difference in administered dose. However, due to the nature of each cell type (as discussed earlier) and the colloidal behaviors of AgNPs in culture media (Ha *et al.* 2018), individual cells within one sample may associate with different numbers of AgNPs and express different levels of cellular responses. The scatter plots of IκBα, STAT1, caspase 7, and cell death levels versus the AgNP uptake of monocytes and dendritic cells (Figures 5 and 6) confirm cell-to-cell variance in AgNP uptake and, consequently, in signaling expression levels. In monocytes and dendritic cells, both NF-κB and JAK-STAT pathways were activated (via degradation of IκBα and phosphorylation of STAT1, respectively) due to the induction of pro-inflammatory and pro-apoptotic signaling as the cellular Ag dose increased. Cell death levels, however, exhibited a direct linear relationship with cellular AgNP uptake. This relationship maintained its linearity in the ^PVP^Ag^10^-treated samples (Figure 5,6B–D), as the cell death level rose accordingly with the incremental increases in cellular Ag dose, from mostly below 10 fg (Figures 5,6B and 5,6C) to over 10 fg, even reaching 100 fg (Figures 5,6D). In addition, caspase 7 cleavage in these samples remained steady regardless of increases in AgNP uptake amounts. This suggests that apoptosis occurred at medium cellular Ag doses (in this case, below 10 fg) and led to cell death at high cellular Ag doses (above 10 fg). On the other hand, the linear relationship between cell death level and AgNP uptake amounts was not very consistent in the ^PVP^Ag^20^-treated samples (Figure 5,6E-G). In this group, the linearity was only maintained when the uptake amounts were below 10 fg. Above this threshold, the uptake amounts remained steady regardless of increases in AgNP uptake. Caspase 7 cleavage in these samples slightly decreased when the cellular Ag dose exceeded 10 fg. This may indicate that above a cellular Ag dose of 10 fg, the phagocytic activities of monocytes and dendritic cells became more efficient and impeded apoptosis signaling, which in turn prevented a rise in cell death level. Similar observations were also reported in other publications, in which apoptosis occurred at a medium cellular dose of quantum dots (QDs), but was impeded by the particle-digestion of cells via autophagy at a higher cellular QD dose, leading to a partial recovery in cell viability at higher doses (Luo *et al.* 2013, Manshian *et al.* 2015). This observation may imply a size-dependent toxicity between the two AgNPs. Because the cell death level of monocytes treated with ^PVP^Ag^10^ NPs still had the potential to rise when the cellular Ag content continued to increase, ^PVP^Ag^10^ NPs should have higher toxicity than ^PVP^Ag^20^ NPs, whose cellular content above 10 fg in mass did not cause an increase in cell death. Such heterogeneous cellular behaviors to subtoxic doses of AgNPs would have gone unnoticed had the response of the larger cell population simply been averaged.

## Conclusion

In this study, we have adapted mass cytometry to investigate the cellular association and cytotoxicity mechanisms of 10 and 20 nm AgNPs in primary human immune cells. Thanks to the high sensitivity of this novel cytometry technique, we could observe the effects of these small AgNPs under non-toxic conditions with low doses (2 and 5 μg/mL) and short exposure time (3 and 6 h). High dimensionality of mass cytometry data allowed us to study the cellular association, cytotoxicity and cell death mechanisms of AgNPs in the heterogeneous immune cell populations. Our results showed that AgNPs were more likely to associate with phagocytic cells like monocytes and dendritic cells, and caused acute toxic effects, but had low affinity to adaptive immune cells such as T cells and B cells, and only caused minor toxicity. In addition to the heterogeneity in associating with different immune cell populations, there was also a great variance between individual cells of each cell type population. Multi-element detection capability of time-of-flight mass cytometry enabled us to quantitatively measure the cellular dose of AgNPs as well as cellular responses simultaneously, so that we could examine the dose-response relationship of cellular AgNPs with cell viability and signaling activities at a single-cell level. We have found that the cell death levels of monocytes and dendritic cells had a linear relationship with the amount of cellular AgNPs. In order to maintain their homeostasis upon the stimuli of AgNPs, monocytes and dendritic cells showed contrasting and complex signaling activities, such as STAT1 phosphorylation to promote inflammation, NF-κB signaling (via degradation of IκBα) to prevent apoptosis and caspase 7 activation to cleave downstream substrates that can eventually lead to apoptosis. These observations suggest that mass cytometry can reveal important information on the interactions between biological cells and NPs, such as heterogeneous association of NPs with different immune cell types, and single-cell level dose-response relationships for cell viability and other signaling activities, which are often hidden in traditional assays at bulk-cell level. We also think that our study will benefit development of single-cell level dose-response models for various NPs and will provide key information for the safe use of nanomaterials for biomedical applications.

## Acknowledgements

Blood samples used in this study were obtained from Hanyang University Hospital (Seoul, Republic of Korea) with the approval of the Institutional Review Board (No. HYUH 2018-09-005-004). Mass cytometry experiments were performed on the Helios CyTOF instrument at the Biotechnology Research Center in POSTECH, Republic of Korea. The authors would like to thank Prof. Young Tae Chang and Dr. Jong Jin Kim (Department of Chemistry, POSTECH, Republic of Korea) for providing access to the CyTOF instrument and helping with its operation. The authors also would like to appreciate Prof. Incheol Shin for his comments on this manuscript.

## Funding

This project has received funding from the Bio & Medical Technology Development Program of the National Research Foundation (NRF) supported by the Ministry of Science & ICT under grant agreement No. 2017M3A9G8084539.

## Author contributions

T.H.Y. designed and supervised research. K.M.H., Z.G., Y. L., Y.-E.K. and J. S. prepared samples. K.M.H. and S.J.K. performed the mass cytometry experiment. K.M.H. and J.S.C. performed data analysis. K.M.H. and T.H.Y. wrote the manuscript.

## Disclosure statement

No potential conflict of interest was reported by the authors.

## Data availability

All the data supporting the findings of this study are available from the corresponding author upon reasonable request.

